# Towards an applied metaecology

**DOI:** 10.1101/422501

**Authors:** Luis Schiesari, Miguel G. Matias, Paulo Inácio Prado, Mathew A. Leibold, Cecile H. Albert, Jennifer G. Howeth, Shawn J. Leroux, Renata Pardini, Tadeu Siqueira, Pedro H.S. Brancalion, Mar Cabeza, Renato Mendes Coutinho, José Alexandre Felizola Diniz-Filho, Bertrand Fournier, Daniel J. G. Lahr, Thomas M. Lewinsohn, Ayana Martins, Carla Morsello, Pedro R. Peres-Neto, Valério D. Pillar, Diego P. Vázquez

## Abstract

The complexity of ecological systems is a major challenge for practitioners and decision-makers who work to avoid, mitigate and manage environmental change. Here, we illustrate how metaecology - the study of spatial interdependencies among ecological systems through fluxes of organisms, energy, and matter - can enhance understanding and improve managing environmental change at multiple spatial scales. We present several case studies illustrating how the framework has leveraged decision-making in conservation, restoration and risk management. Nevertheless, an explicit incorporation of metaecology is still uncommon in the applied ecology literature, and in action guidelines addressing environmental change. This is unfortunate because the many facets of environmental change can be framed as modifying spatial context, connectedness and dominant regulating processes - the defining features of metaecological systems. Narrowing the gap between theory and practice will require incorporating system-specific realism in otherwise predominantly conceptual studies, as well as deliberately studying scenarios of environmental change.

---

For centuries humans have destroyed, modified, overharvested and polluted ecosystems, leading to irreversible environmental damages such as species extinctions and the loss of key ecosystem services. Avoiding, mitigating, and managing environmental change is therefore one of the greatest challenges for mankind in the 21^st^ century (MEA 2005a). So far, the search for solutions has revealed two important lessons. First, that the ecological processes involved are rarely if ever purely local because ecological systems are open and connected through the flow of organisms, energy and matter. Second, that managing environmental change and related consequences frequently requires addressing processes at greater spatial scales than those where observed changes took place. Yet, we are still in need of a conceptual framework bridging the empirical knowledge about the processes underlying biodiversity and ecosystem function with environmental management, while adequately addressing issues of scale. Ecosystem-based management advanced as a holistic approach to address many of the failures of traditional natural resource management (Slocombe 1998) but is not adequately linked with spatial ecological theory. Macrosystems ecology has emerged as a framework for considering human-ecological interactions at continental extents (Heffernan et al. 2014), but doesn’t address how many management decisions and responses that occur at local or landscape extents apply. Here, we propose the use of ′metaecology′ to address critical environmental issues by providing a more holistic multi-scale framework to understand and predict the ecological consequences of environmental change. Throughout this paper, we use ‘metaecology’ as a grouping term for metapopulation (a set of local populations of a single species that are linked by dispersal; Levins 1969), metacommunity (a set of local communities that are linked by dispersal of multiple interacting species; Wilson 1992) and metaecosystem ecology (a set of local ecosystems that are linked by the flow of organisms, energy or matter; Loreau et al.2003), and define it more broadly as the study of interdependencies among ecological systems through fluxes of organisms, energy, or matter in space (Figure 1, Box 1).

We first provide a series of case studies illustrating the use of metaecology to address management or conservation problems. Subsequently, we present a literature review to make a critical assessment of the use of metaecological concepts in applied ecological studies and in recommended management responses to drivers of environmental change. We then discuss how environmental change can be conceptualized within the framework of metaecology, and end by highlighting a series of challenges and future directions underlying environmental management at local and regional scales.

**Figure 1.**
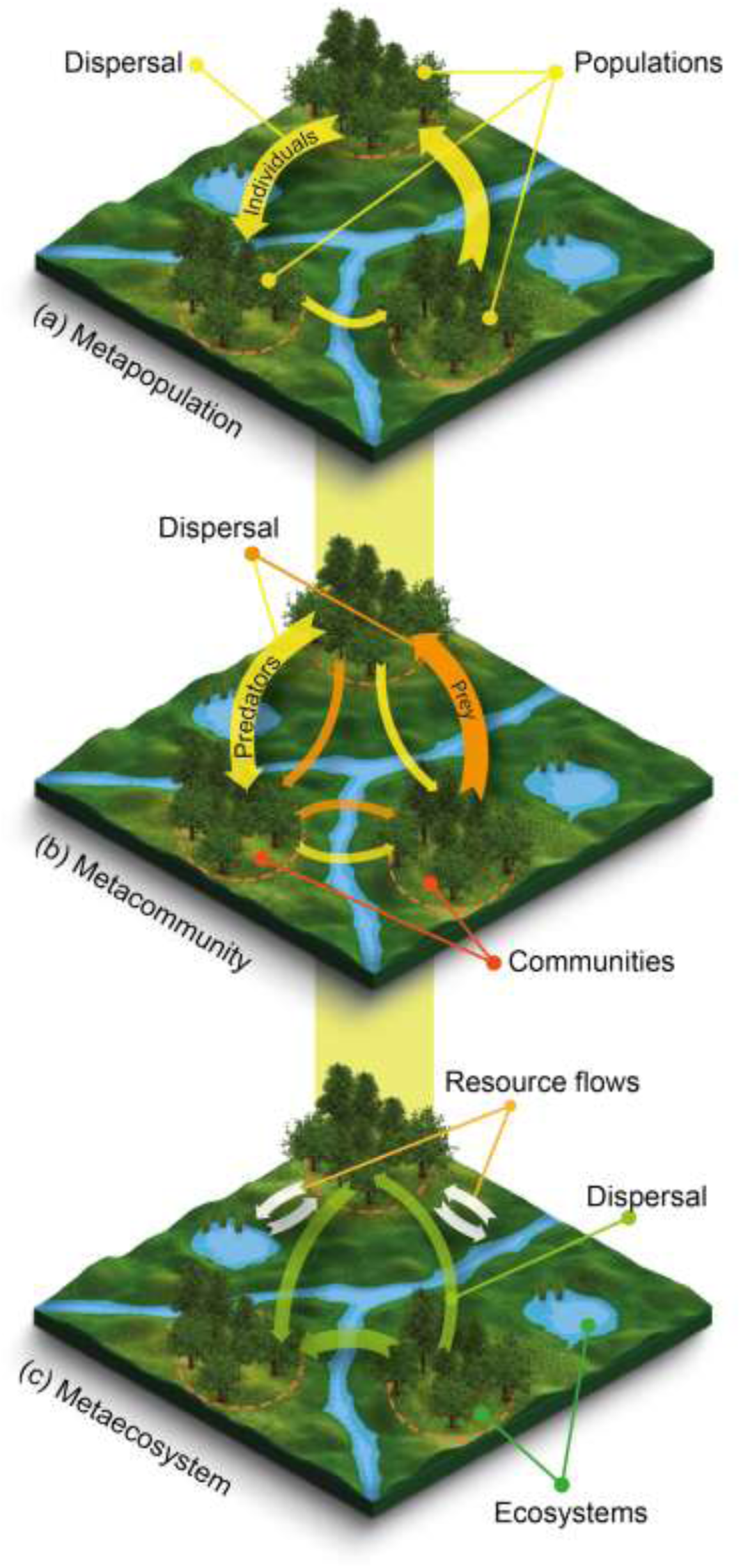
Schematic representation of metaecological entities: a. a metapopulation, b. a metacommunity, and c. a metaecosystem. In each landscape, the orange dashed circle indicates local populations, communities or ecosystems in forest patches connected through the flow of organisms (colored arrows) and resources (white arrows). Note that flow rate (arrow thickness) may vary across the landscape. Even though metacommunities are defined as sets of communities linked by the dispersal of individuals of multiple interacting species, for the sake of simplicity much of what ecologists have studied as metacommunities are actually sets of assemblages of species within the same trophic level (not shown). Resource flows represented in the metaecosystem could be forest patches contributing leaf litter to streams and lakes, in turn contributing emerging insects for birds foraging in forest patches.

## Addressing real-world problems with metaecology

The link between metaecology and applied ecology is not entirely new and, in fact, recent developments of this framework have been prompted by the need to address challenges in conservation biology and environmental management (Bengtsson 2009). The five case studies in this section illustrate how metaecological approaches can be used to support managerial decisions at different levels of biological organization.

### Case study 1: Incorporating population connectedness to design protected areas

Metapopulation ecology has been instrumental in developing optimization procedures to design protected area networks for the conservation of endangered species (Hanski 2004). The false heath fritillary butterfly (*Melitaea diamina*) historically benefited from traditional agriculture in Finland because small-scale tilling and mowing favours its host plant, the pioneer valerian (*Valeriana sambucifolia*) (Figure 2). However, due to natural succession and land use change, the species range plummeted to two small regions where it persisted as a metapopulation in a dynamic network of suitable habitat patches connected by dispersal (Cabeza 2003). Metapopulation models and dispersal studies identified priority management sites, and motivated the development of applied conservation software for identifying optimal sets of conservation sites accounting for cost (Moilanen and Cabeza 2002) and spatial dynamics (Cabeza 2003). This successful application of metaecology highlights how the framework effectively confronts the challenges of developing optimal conservation strategies when suitable habitat patches shift in space and time, and eventually led to an evaluation of the spatiotemporal fit of Finnish institutions involved in the species protection (Fabritius et al. 2017). Spatial structure, connectedness, turnover and quality of local patches are now continuously monitored by local conservation authorities in Finland, and used to revise alternative strategies for urban planning. These have been pivotal in guaranteeing the persistence of *Melitaea diamina* in the country.

**Figure 2.**
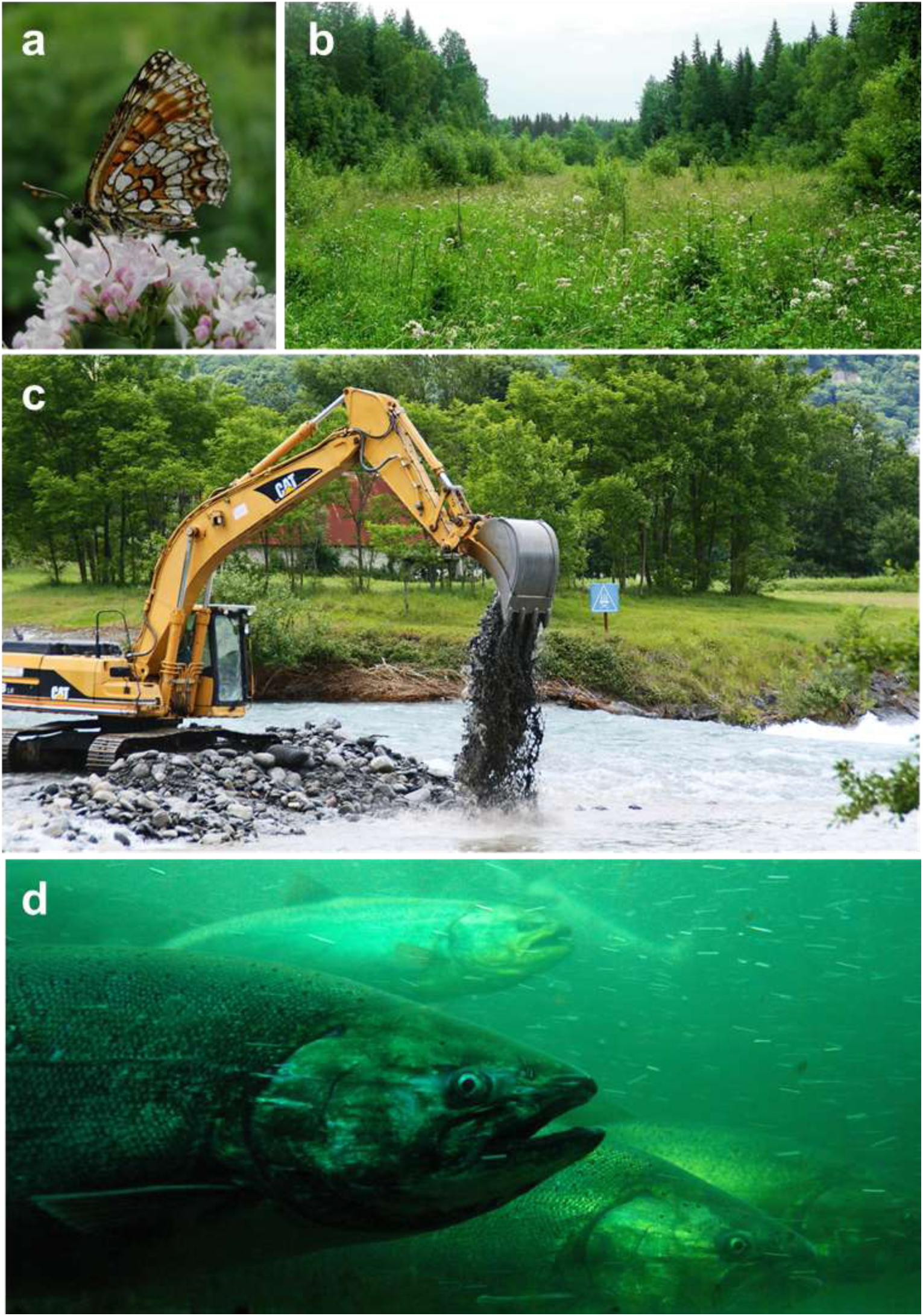
**a-d**. Selected case studies. **a,b**. Metapopulation studies of the endangered false heath fritillary butterfly (*Melitaea diamina*) prompted the development of algorithms for identifying optimal sets of conservation sites accounting for both cost and spatiotemporal dynamics. Accounting for spatiotemporal dynamics was a necessity because the butterfly obligatory host is a pioneer plant (the valerian *Valeriana sambucifolia*), which distribution shifts as a function of forest succession and land use change. **c**. Stream restoration historically focused on the manipulation of local habitat heterogeneity (e.g. reconfiguration of the channel, addition of logs, boulders and gravel), implicitly assuming that species would spontaneously recolonize. It was more recently found that, because contemporary dispersal among streams is very rare, community recovery requires consideration of proximity and active pathways to colonization sources, or deliberate stocking. **d**. Pacific salmon (such as the chinook salmon *Onchorhyncus tshawytscha)* can accumulate in its biomass large quantities of biomagnifiying metals and persistent organic pollutants, and effectively transfer them to sometimes distant ecosystems via upstream migrations and death. Figures courtesy of Kale Meller (a, b), Herbythyme (c; GFDL/Creative Commons; image cropped) and Josh Larios (d; Creative Commons).

### Case study 2: Designing habitat networks to the long-term sustainability of multispecies landscapes

Beyond a single endangered, umbrella or flagship species, protected area networks should satisfy the requirements of multiple species with contrasting life histories and movement ecologies. Albert et al. (2017) applied the framework of metaecology to prioritize forest remnants for the long-term conservation of fourteen vertebrate species ranging from salamanders to bears in the St. Lawrence lowlands surrounding Montreal, Canada. For conservation to be effective in the long term, the authors accounted for future land use and climate change, and their uncertainties. The solution comprised a combination of well-connected, large forest patches favoring the short-range connectedness that is required for the persistence of metapopulations within the network, with corridors of smaller stepping-stone patches promoting long-range connectivity that facilitates climate-driven range shifts across adjacent networks. The design of such habitat network was a direct request from the Ministry of Sustainable Development, Environment and Parks of the Quebec government and a strong relationship with local stakeholders ensured that these results are now being concretely used in the design of Montreal’s greenbelt.

### Case study 3: Preserving plant–pollinator interactions in agricultural landscapes

While the previous case study considered the individual responses of multiple, non-interacting species (i.e., a collection of metapopulations), ecologists and practitioners are sometimes interested in managing species interactions. This is the case of plant-pollinator mutualisms. Because many crop species depend heavily on animal pollination to produce fruits and seeds (Garibaldi et al. 2013), the increased loss of remnants of natural vegetation in intensively managed agricultural landscapes poses a serious threat not only to biodiversity conservation but also to agricultural yields (Bengtsson 2009). Metaecological approaches have provided explicit recommendations as to how to manage habitat structure, restore non-agricultural habitat, and/or enhance connectedness between agricultural fields and surrounding natural or seminatural habitats. For example, restoring hedgerows in agricultural landscapes can improve the richness and abundance of pollinators which, by flowing into interconnected fields, may maintain and enhance the yield, quality and diversity of agricultural products (Morandin and Kremen 2013). Reducing the impacts of agricultural intensification require taking into account interaction at local scales but also how different habitats are interconnected and distributed across space.

### Case study 4: The role of connectedness in restoring degraded communities

Stream restoration has historically focused on the manipulation of local habitat heterogeneity (e.g. reconfiguration of the channel, addition of logs, boulders and gravel; Figure 2), implicitly assuming that individuals of different species would spontaneously recolonize restored reaches. However, many of these restoration projects failed because target species were either absent in the regional species pool or unable to disperse to restored sites (Palmer et al. 2014). Considering dispersal and metacommunity dynamics is critical in river networks since isolated tributaries receive few immigrants from the main river corridor and most freshwater organisms rarely disperse overland among streams or catchments (Hughes 2007). Thus, communities in isolated headwaters are predicted to be more vulnerable to environmental change than communities inhabiting well-connected downstream branches. Swan and Brown (2017) tested this idea using metacommunity ecology to study how restoration efforts fared in headwater *versus* mainstem sites. Their study clearly demonstrates that restoration efforts are context-dependent, enhancing biodiversity and stability of ecological communities in more isolated headwater sites, where local processes play a major role, while similar efforts in well-connected mainstems are largely ineffective because regional processes dominate. Even so, successful restoration of headwaters requires recolonization and maintenance of viable populations in restored stream sites through proximity to colonization sources (Parkyn and Smith 2011) or deliberate introductions (Stranko et al. 2012). This case study demonstrates that building management actions considering spatial structure and dispersal among target streams can increase the success of restoration activities.

### Case study 5: Animal-mediated flows of pollutants among ecosystems

Spatially separated ecosystems may be connected to each other by the flow of energy and materials – including chemical contaminants. Throughout the 20th century, the Laurentian Great Lakes and their basins were subject to heavy discharges of persistent organic pollutants (POPs) including PCBs (polychlorinated biphenyls), PBDEs (polybrominated diphenyl ethers) and DDE (dichlorodiphenyldichloroethylene), which were used in household and industrial products such as electrical transformers, flame retardants, and pesticides. Known primary routes of contaminant dispersal from release sites included volatilization and atmospheric deposition, as well as runoff and downstream transport of water and eroded sediment. Not considered was the spatial redistribution of these POPs by the regular stocking of salmonids for recreational and commercial fisheries (745 million fish stocked in the Great Lakes since 1967; Crawford 2001). PCBs, PBDEs and DDE biomagnify along food chains and predatory pacific salmon such as *Oncorhynchus tshawytscha* (Figure 2) can accumulate high POP concentrations in their biomass and transfer them to tributaries via upstream spawning migrations and death (Janetski et al. 2002). Decomposition and consumption of salmon carcasses facilitate POP flow through local food webs and contaminate resident brook trout, a species targeted by recreational anglers and not regularly assessed for contaminant levels nor included in health consumption advisories (Janetski et al. 2002). Interestingly, whereas an increase in connectedness was a target for environmental managers in all case studies above, here the costs and benefits of increased connectedness must be weighed: considering that hundreds of dams are being removed as part of the Great Lakes Restoration Initiative, we can anticipate salmon-mediated contamination of newly accessible upstream tributaries to increase in this metaecosystem. More generally this case study illustrates that the movement of materials across ecosystems (from lake to tributaries) may have important consequences for human welfare.

### Beyond selected case studies: scale-restricted trends in applied ecology

These five case studies illustrate how metaecology can help address important environmental issues more effectively across scales and at multiple levels of biological organization, but the wider use of metaecological approaches is still relatively rare in applied ecology. We conducted a formal bibliometric analysis of the applied ecology peer-reviewed literature to (i) quantify the pervasiveness of metacological concepts and (ii) assess whether explicit metaecological frameworks are being used in studies of environmental change, based on their use of terminology, study design, and literature cited. Our analysis shows that very few of the surveyed papers used or referred explicitly to a metaecological approach (Box 2). Additionally, we analysed published guidelines for mitigating or managing the risk of the key drivers of environmental change, which generally revealed a widespread focus on local scale processes, and a lack of focus on spatial context (spatial hierarchy and heterogeneity) and connectedness (Box 3). Despite scientific studies (Box 2) and management actions (Box 3) seldom explicitly considered metaecology, they frequently incorporated two or all three of its key features. Connectedness (i.e. flow) was the least considered feature in the studies reviewed here, even though it is one of the key aspects to be addressed in management actions. Among drivers of environmental change, studies on land use change were more likely to incorporate connectedness while studies addressing pollution and overharvesting rarely acknowledged it.

## Framing environmental change within metaecology

More generally, we argue that metaecology provides a valuable framework for understanding and managing the consequences of environmental change because of its impacts on all three defining features of interdependent ecological systems.

a. <u>Environmental change alters the intensity and/or direction of natural flows of organisms, energy and materials across landscapes.</u> One of the most important contributions of metaecology is the recognition that connectedness strongly influences ecological dynamics (Box 1). Modifying connectedness is one pervasive consequence of environmental change. For example environmental change can decrease connectedness – such as in river damming (Haxton and Findlay 2008) and habitat fragmentation (Fahrig 2003) – or increase connectedness – such as in the intercontinental transfer of marine organisms in ballast water (Seebens et al. 2013) and the global redistribution of pollutants in the atmosphere (Franklin 2006). Metaecology argues that the consequences of changing connectedness is frequently non-linear (e.g. Mouquet and Loreau 2003, Leibold et al. 2017) so that simple linear expectations are not likely to apply in general.

b. <u>Environmental change modifies the spatial context, from local to regional scales.</u>

Environmental change generates novel, spatially structured stress signatures that are capable of reinforcing, modulating or overriding the spatial structure that prevailed under natural conditions. Like other factors of ecological relevance (Box 1), the spatial structure of anthropogenic drivers of environmental change can also be described in terms of spatial heterogeneity and spatial hierarchy. The spatial structure of environmental change is evident, for example, by the non-random distribution of tropical deforestation hotspots and of entrance points of biological invasions, like ports and harbors. This heterogeneous distribution of drivers of environmental change implies that the magnitude of stress that any given ecological entity is exposed to is predictable by its distance to the driver′s source, but also by the spatial extent of influence of the driver itself. Some drivers have a very local extent of influence (i.e. influence only ecological systems that are in close proximity) whereas others may have regional or even global ranges of influence (Figure 3). Spatial hierarchy in turn means that drivers can also scale up or down. Air pollution, for example, originates from local emission sources and at this scale influences biodiversity and ecosystem function by acidification, ecotoxicity and CO_2_ enrichment. However, a multitude of local sources of air pollution leads to cumulative or synergistic changes at planetary scale (i.e., greenhouse effects) that manifest as global climate change (Turner et al. 1990). Metaecology provides conceptual ways to understand these scale dependent effects and link them to the response of ecosystems.

**Figure 3.**
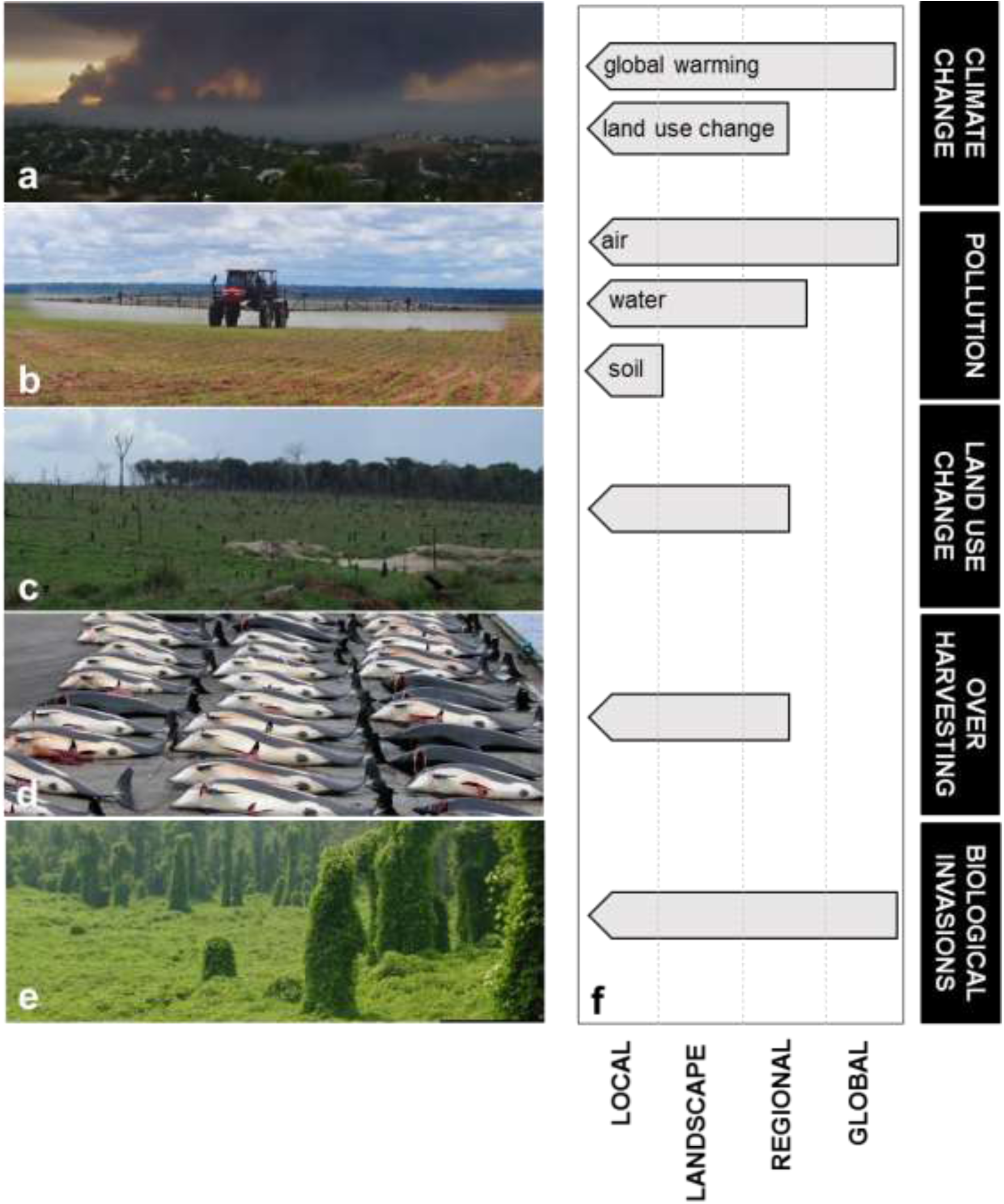
**a-e**. Main anthropogenic drivers of environmental change. **a**. Climate change (greenhouse gas emission by wild fires in California, USA). **b**. Pollution (pesticide application in Southern Amazon, Brazil). **c**. Land use change (conversion of rainforest into pasture in Southern Amazon, Brazil). **d**. Overharvesting (whaling of Atlantic White-sided Dolphins in the Faroe Islands, Denmark) **e**. Biological invasions (the vine kudzu growing over native vegetation, eastern USA). **f**. The variable spatial extent of drivers of environmental change on local ecological entities (population, community or ecosystem). Using pollution as an example, a local ecological entity could be under the influence of local deposition of solid waste, of regional release of untreated wastewater in the drainage network upstream, or of global atmospheric pollution of carbon dioxide, nitrate, dust, and persistent organic pollutants. The typical maximum spatial extent of influence of each driver on a local ecological entity is depicted. Two different mechanisms driving ′climate change′ are represented, one of global (greenhouse effect) and one of landscape-to-regional extent of influence (land use change). Figures a, d and e courtesy of Nerval (Public Domain in Wikipedia Commons), Erik Christensen (GFDL/Creative Commons; image cropped) and Kerry Britton (USDA Forest Service, Bugwood.org).

c. <u>Environmental change disrupts ecological processes from local to regional scales.</u> Drivers of environmental change operating at small spatial scales are likely to affect local species diversity, migration rates, and the environment. However, spatial heterogeneity in these drivers (‘b’ above) can disrupt trophic interactions and ecosystem properties by modifying the level of connectedness between habitats (‘a’ above), as determined by the contrasting dispersal distances and rates of species in different taxonomic and functional groups (e.g. predators and prey) and energy flows (Sitters et al. 2015). These consequences will have different outcomes from local to regional scales. For example, loss of connectedness affects species composition at small scales but not at larger scales (Brose and Hildebrand 2016). Understanding these scale dependencies in response to environmental change is crucial when considering management actions which may be ineffective if particular trophic levels are dependent on biotic or abiotic resources isolated in distant patches or unavailable in the landscape as a consequence of environmental change (Montoya et al. 2012). To effectively sustain or increase biodiversity at multiple scales, management should target species-rich sites supporting remnants of the original native assemblage that maximize ecosystem function, and that can be spatially well-connected to other similar sites in the region.

## Recent advances and future challenges in applied metaecology

Even though it has already provided key insights about the spatial dynamics of ecological systems, it is important to realize that metaecology is still a rapidly developing field. In particular there a numerous recent and ongoing conceptual, theoretical, and methodological developments that will increasingly narrow the gap between theory and practice in metaecology. Initial approaches to metaecology primarily emphasized spatially implicit theories (that is, where actual spatial structure did not matter) in search of generality. This made the connection with management difficult because management almost always requires a spatially explicit context. However more recent work in metaecology theory is moving in this direction (e.g. Economo 2011, Marleau et al. 2014), as well as in developing modelling toolkits for predicting species responses to environmental heterogeneity and change (Keyel et al. 2016). Likewise, metaecology is moving from simplified general assessments of the effects of connectedness towards more realistic incorporation of variation in flow rates and flow modalities among species, materials and landscapes (e.g. Bohonak and Jenkins 2003, Riibak et al. 2014, Fournier et al. 2016). These trends will undoubtedly continue and make metaecology more flexible and realistic in ways that will continue to improve understanding and facilitate how we understand complex socio-ecological systems.

Although we have highlighted how applied ecologists could benefit from better incorporating metaecology, many of the important environmental issues also present important challenges that could greatly strengthen metaecology by highlighting topics that haven’t yet been particularly well incorporated. Some of these include topics related to:

a. *<u>Responding to climate change</u>*. Under a changing climate, it is predicted that species will be redistributed across space through natural or human-mediated geographic range shifts following climate-driven alterations to local habitat suitability (Hoegh-Guldberg et al. 2008, Pecl et al. 2017). These shifts in species ranges will modify the composition and functional diversity of regional species pools, leading to novel species assemblages (Moritz and Agudo 2013). Metaecology offers a spatial framework that has the capacity to predict changes in species diversity across multi-scale species diversity and ecosystem function as a consequence of these large-scale changes in species distributions (e.g. Norberg et al. 2012).
b. *<u>Predicting, avoiding and controlling biological invasions.</u>* As a product of geographic range shifts mostly in response to climate change and human-mediated dispersal, biological invasions are an increasing threat to the biotic integrity of ecosystems as global homogenization ensues (Baiser et al. 2012). Predicting, monitoring, and controlling the spread and impact of invasive species proves to be a major ecological and economic challenge (Lodge et al. 2016). Metaecology combines metapopulation models of individual invasive species with metacommunity and metaecosystem models. Considering how metacommunity and metaecosystem connectivity can prevent or facilitate the establishment and regional spread of invasive species will be critical for conservation and management (Howeth 2017).
c. *<u>Improving land use planning</u>*. Habitat patches vary greatly in size, structure, diversity and connectedness, and recent work indicates that certain patches play disproportionately larger roles in maintaining biodiversity or other community/ecosystem attributes in a landscape (Tews et al. 2004, Economo 2011, Mouquet et al. 2013). Acknowledging and identifying such ′keystone habitats′ and ′keystone communities′ is at the core of prioritization efforts in reserve design, landscape management and restoration. In addition, landscapes are not only heterogeneous but also dynamic due to habitat fragmentation and the shifting nature of managed landscapes. Because metaecology has so far mostly focused on static landscapes (Sferra et al. 2017), addressing how changing environmental conditions and landscapes alter metacommunity and metaecosystem dynamics is a major research goal for metaecology that can benefit from the insights gained from applied studies (see also van Teefelen 2012).
d. *<u>Controlling pollution and its consequences.</u>* Quantifying and predicting the dispersal of contaminants has been a central goal of contaminant fate models, but controlling connectedness is not usually an important component in pollution management. Abiotic processes have usually been assumed to dominate contaminant transport and processing but organismal movement can directly or indirectly contribute to contaminant fate since individuals act as contaminant reservoirs and/or processors. Because these roles can be partly inferred from contaminant properties and organismal traits, integrating contaminant properties, organismal traits, community structure and connectedness may help us anticipate and better manage the spatial redistribution of contaminants, as well as the environmental context by which ecosystems turn from contaminant sinks to sources and vice-versa (Schiesari et al. 2017).

## Conclusions

This article demonstrates that metaecology provides many important insights about the dynamics of populations, communities and ecosystems in a changing environment, which have directly contributed to conservation, restoration, and risk management actions. There are, nevertheless, a variety of questions that require further theoretical, methodological and/or empirical developments if metaecology is to underpin environmental management, as the rapid growth of the field has been largely conceptual. Clearly, it is the iteration between scientists, practitioners and decision-makers that will provide the impetus for this development. We propose a rule-of-thumb to determine whether a metaecological framework is required, and thus demand more explicit recommendations from scientists, based on two questions: (i) does environmental change vary across space and/or modify the spatial distribution of natural ecosystems and resources? and (ii) does this change alter the intensity and direction of natural flows (organisms, energy and materials) across landscapes? If the answers are both ‘yes’, then we strongly advocate that a metaecological framework should be employed for three reasons. First, metaecology offers a strong theoretical and analytical foundation to detect and model responses to environmental change. Second, metaecology – through a deep understanding of connectedness and spatial context – can improve management from local to regional scales. Finally, metaecology has the potential to identify mismatches between the scales at which ecological processes are operating and those at which political decision-making and environmental management occur. Closing this gap is essential to better assess environmental problems and to find politically and ecologically sustainable solutions (Cash et al. 2006).

## Acknowledgments

This article is a product of the ′Applied Metaecology Workshop′, organized by LS, PIP & MAL in Ilhabela, Brazil, from March 13-17, 2016 and funded by FAPESP (São Paulo Research Foundation; grant 2015/17984-9). We thank Melina Leite and the Graduate Program in Ecology of the University of São Paulo for logistical support. We thank FAPESP (grants 2014/10470-7 to AM, 2013/04585-3 to DL, 2013/50424-1 to TS and 2015/18790-3 to LS), CNPq (Productivity Fellowships 301656/2011-8 to JAFDF, 308205/2014-6 to RP, 306183/2014-5 to PIP and 307689/2014-0 to VDP), the National Science Foundation (DEB 1645137 to JGH), the Natural Sciences and Engineering Council of Canada (SJL, PPN), and the Academy of Finland (grants 257686 and 292765 to MC) for support. This work contributes to the Labex OT-Med (no. ANR-11-LABX-0061), funded by the French government through the A*MIDEX project (no. ANR-11-IDEX-0001-02). We thank Jan Bengtsson for constructive criticism on a previous version of this manuscript.

## Box 1. The three key defining features of metaecology

### Spatial structure in the environment

The properties of a hypothetical environment that is completely homogeneous and well-mixed, and therefore lacking spatial structure, are usually unaffected by the movement of organisms and materials. Otherwise, spatial structure in the environment can be an important regulator of the dynamics of populations, communities, and ecosystems. Spatial structure has two main components: spatial hierarchy and spatial heterogeneity. Spatial hierarchy refers to the different levels of spatial organization of ecological systems. In the simplest view, this hierarchy involves a small scale (locality), where non-spatial processes may be particularly important (e.g. demography and species interactions), and a larger scale (region), where the dominant processes might differ (e.g. dispersal and colonization history). Spatial heterogeneity refers to the degree by which the spatial distribution of an ecological factor (such as temperature, moisture or nutrient availability) or process (such as predation pressure) varies over space. Heterogeneity and its spatial distribution can be important in affecting metapopulations (Hanski 1994), metacommunities (Logue et al. 2011) and metaecosystems (Gounand et al. 2018). To date, metaecology has mostly addressed spatial structure by thinking of discrete habitat patches separated by an inhospitable matrix that limits dispersal, but ongoing work is likely to change this simplistic framework (e.g. Chase and Knight 2013, Garzon-Lopez et al 2014, Munkenmuller et al. 2014).

### Connectedness

The movement of organisms and materials results from the interaction between their dispersal and flow rates, respectively, and the spatial organization of the environment. This interaction determines spatial connectedness, and it is this connectedness that drives important features of the overall system at both local and regional scales. For some processes, the dynamics can be thought to be ‘flow limited’ because colonists are not immediately available to establish populations, or because spatial flows of key materials are limiting to material cycles. For other processes the dynamics can be thought to be subject to ‘flow excess’ because the dispersal of colonists or the flow of materials from other locations can be so high as to overwhelm local dynamics. For example, flow excess can support source-sink population relations (Pulliam et al. 1988) or provide spatial subsidies of materials (Polis et al. 2004) in the extreme leading to spatial homogenization. Intermediate connectedness wherein colonization is (nearly) immediate and material flows are not limiting to material cycles nor are they so large as to lead to spatial homogenization have their own dynamic; one that leads to closer correspondence between local abiotic environmental conditions and biotic features of communities such as species composition and internally balanced material dynamics. Of course, this means that different processes can have different effective scales depending on the connectedness of the important agents involved.

### Localized underlying ecological mechanisms

Finally, metaecology recognizes that critical components of the dynamics of ecological systems are sensitive to, and occur in, spatially restricted locations. For individual species this corresponds to the classic meaning of a ‘local population’ and can be related to the dispersal of individuals. For communities and ecosystems, the spatial definition of ‘community’ or ‘ecosystem’ is less concrete, and indeed served as a long discussed topic (Wiens 1989). Nevertheless, key elements of the dynamics of populations (e.g. demography and vital rates), communities (e.g. species interactions and responses to local environmental conditions), and ecosystems (e.g. element cycling and stoichiometric linkages among material fluxes) occur over strongly localized extents. The overall properties of the system emerge from the way these localized mechanisms interact with the spatial processes in metaecology.

## Box 2. How often have applied ecologists employed a metaecological framework?

We conducted an initial bibliometric analysis to assess how often metaecology is employed in the applied ecology peer-reviewed literature (as indicated by the usage of the terms ′metapopulation′, ′metacommunity′ or ′metaecosystem′ in title, abstract or keywords; eight journals; see WebPanel 1 for Methodology). We found 672 references for ′metapopulation′, 51 for ′metacommunity′ and only 2 for ′metaecosystem′. This reflects, at least in part, the chronology of the field: the term ′metapopulation′ was coined in 1969 (Levins 1969); ′metacommunity′ in 1992 (Wilson 1992); and ′metaecosystem′ in 2003 (Loreau et al. 2003). We conclude that, apart from metapopulation studies, metaecology has been rarely employed in applied ecology – at least not formally.

We then conducted a second bibliometric analysis to assess whether published research investigating specific drivers of environmental change explicitly employed the metaecological framework based on their use of terminology, study design, and literature cited. We randomly selected 25 recent (2011-2016) papers for each of the five main drivers of environmental change using search words relative to biological invasions, overharvesting, land use change, pollution and climate change in the title, abstract and keywords. Each paper was randomly assigned to one of six pairs of evaluators (see Supplementary Materials for Methodology).

Only 3% of the articles referred to metaecology (i.e., used the words ′metapopulation′, ′metacommunity′ or ′metaecosystem′) and only 8% cited any literature recognizable as pertaining to the field (Table 1). Nevertheless, many of the articles included in the study design two or more of the key features of metaecology. Most studies (83-86%) considered multiple locations (ranges represent among-evaluator variation in interpreting the selected literature). This was frequently due to a need for spatial replication; when we restricted our analysis to cases that considered the hierarchy of spatial structure, i.e., where the collective properties of multiple locations where reported (e.g., beta and/or gamma diversities in assemblage studies), values dropped to 23-32%. Key ecological processes at the local scale such as demography, single species responses to the abiotic environment, or species interactions were commonly reported (56-64%), and a smaller fraction directly or indirectly addressed the flow of individuals, matter or energy (34-41%).

**Table 1.**
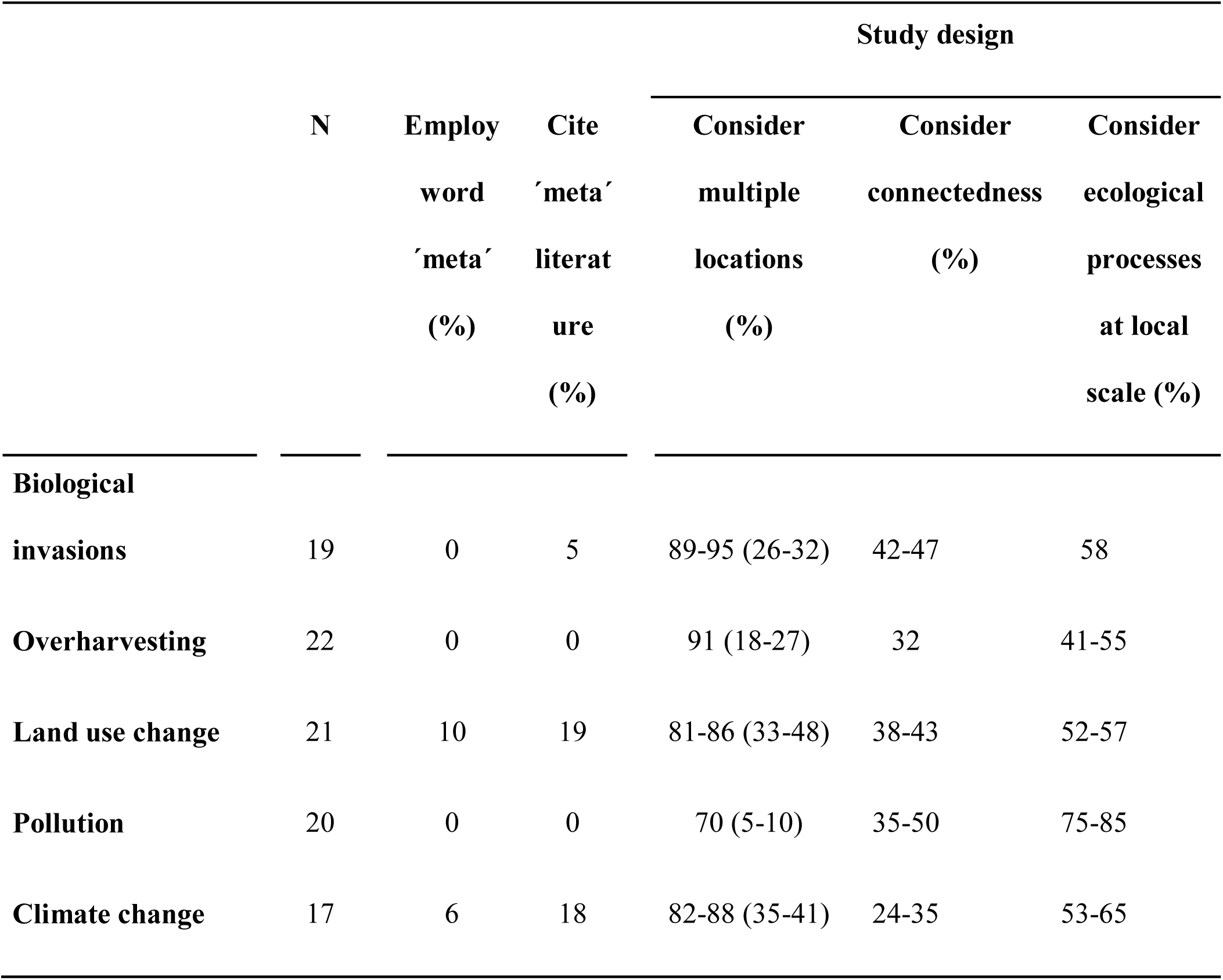
Bibliometric analysis regarding the employment of elements of the metaecological framework (as judged by terminology, literature cited, and study design) in a random selection of recent [2011-2016] articles about the five most important drivers of environmental change. All values are percentages of the articles. Among-evaluator variation in interpreting the selected literature is presented as ranges. In the column ′Consider multiple locations′, values inside the parentheses indicate articles with a hierarchical consideration of spatial structure, i.e., where the collective properties of multiple locations where reported (e.g., beta and/or gamma diversities in assemblage studies). See WebPanel 1 for detailed Methodology.

## Box 3. How often have practitioners employed a metaecological framework?

Perhaps most appropriately for practitioners, we evaluated the correspondence between recommended responses (i.e. actions) to drivers of environmental change and the key defining features of metaecology. We analyzed guidelines and/or reviews of responses to biological invasions, overharvesting, land use change, pollution and climate change (IUCN 2000, MEA 2005b,c,d,e,f, Grafton et al. 2010) and organized such responses as either pertaining to the management of regional spatial structure, connectedness, or local processes (Table 2).

**Table 2.**
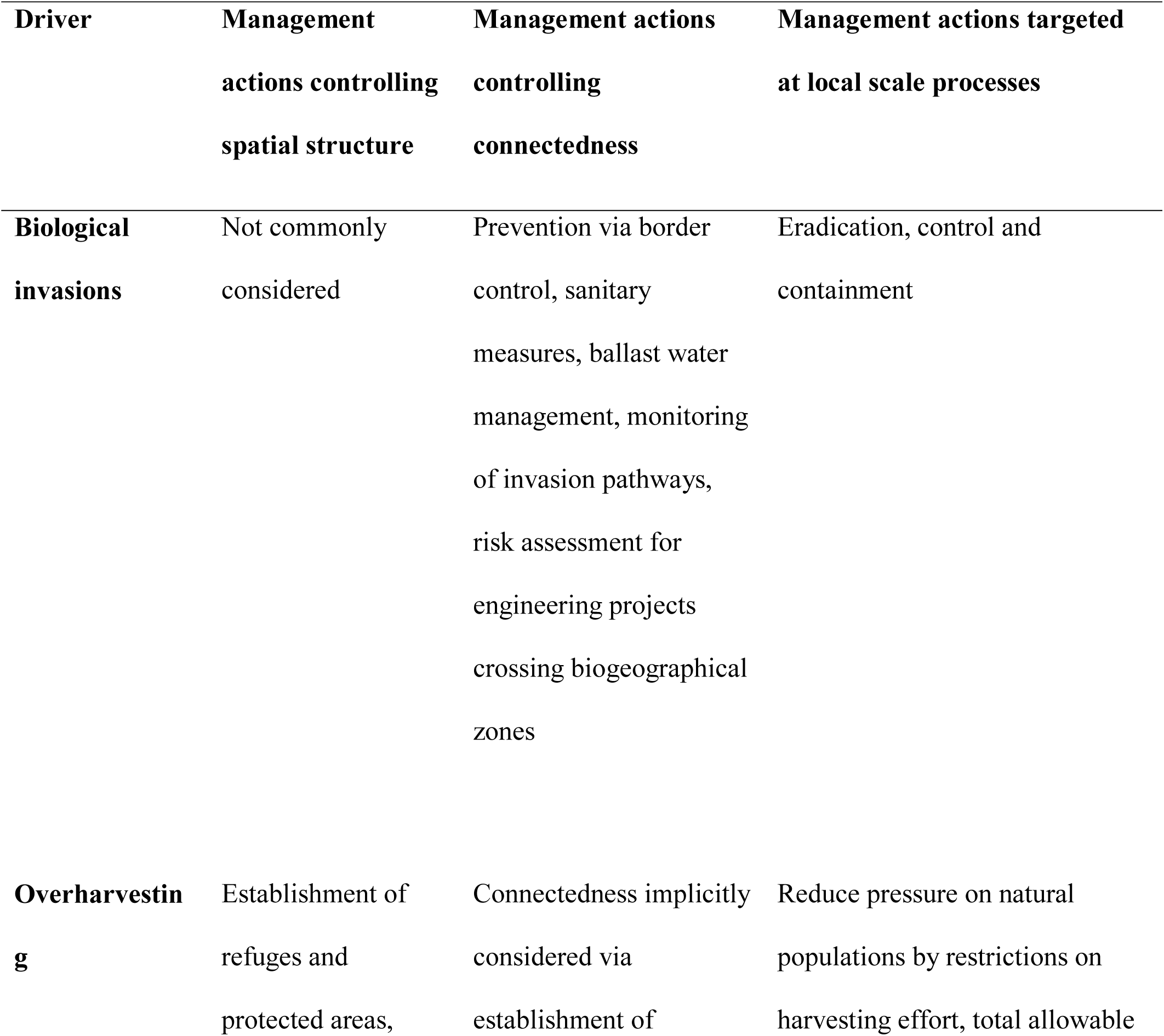
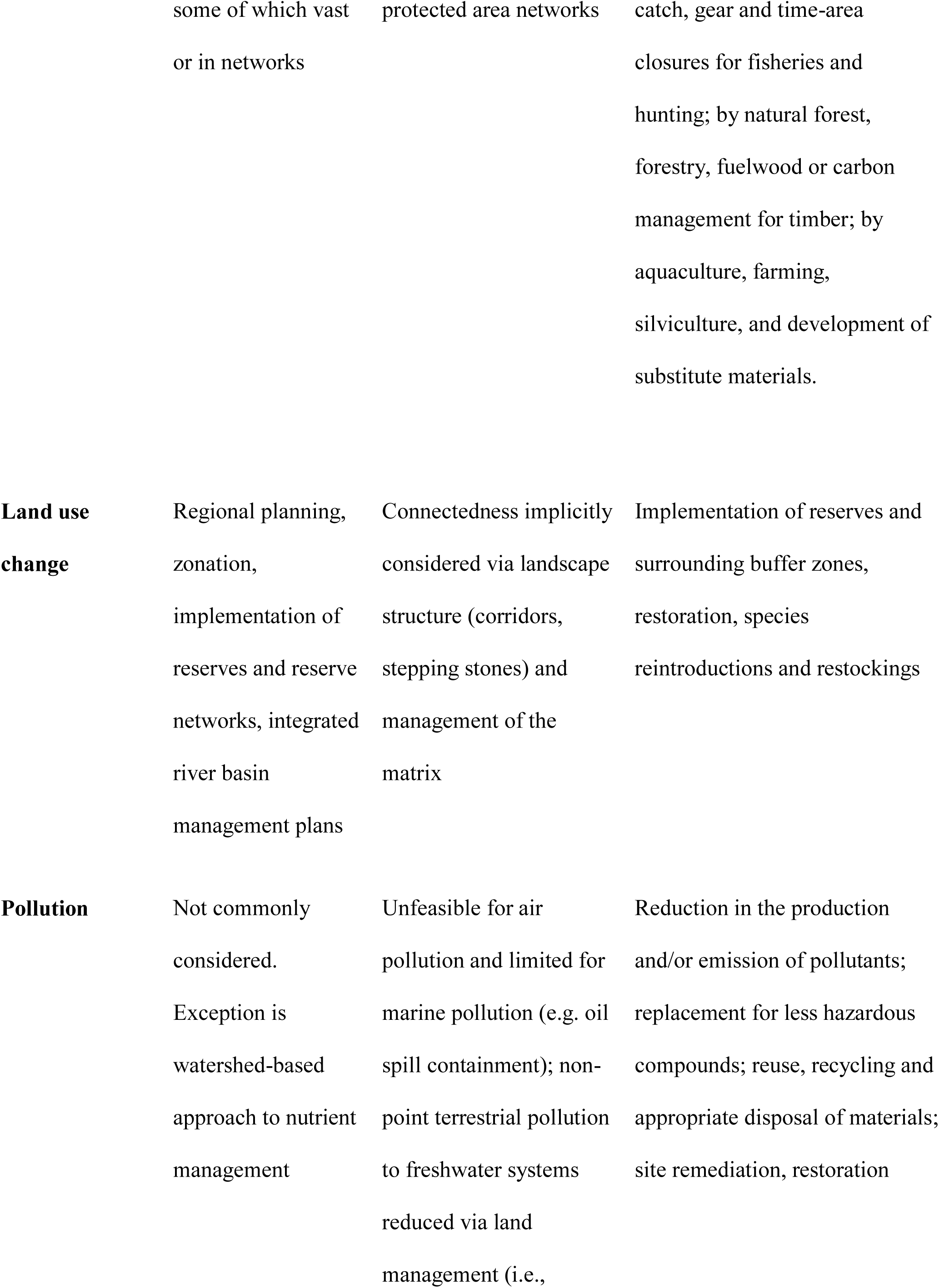
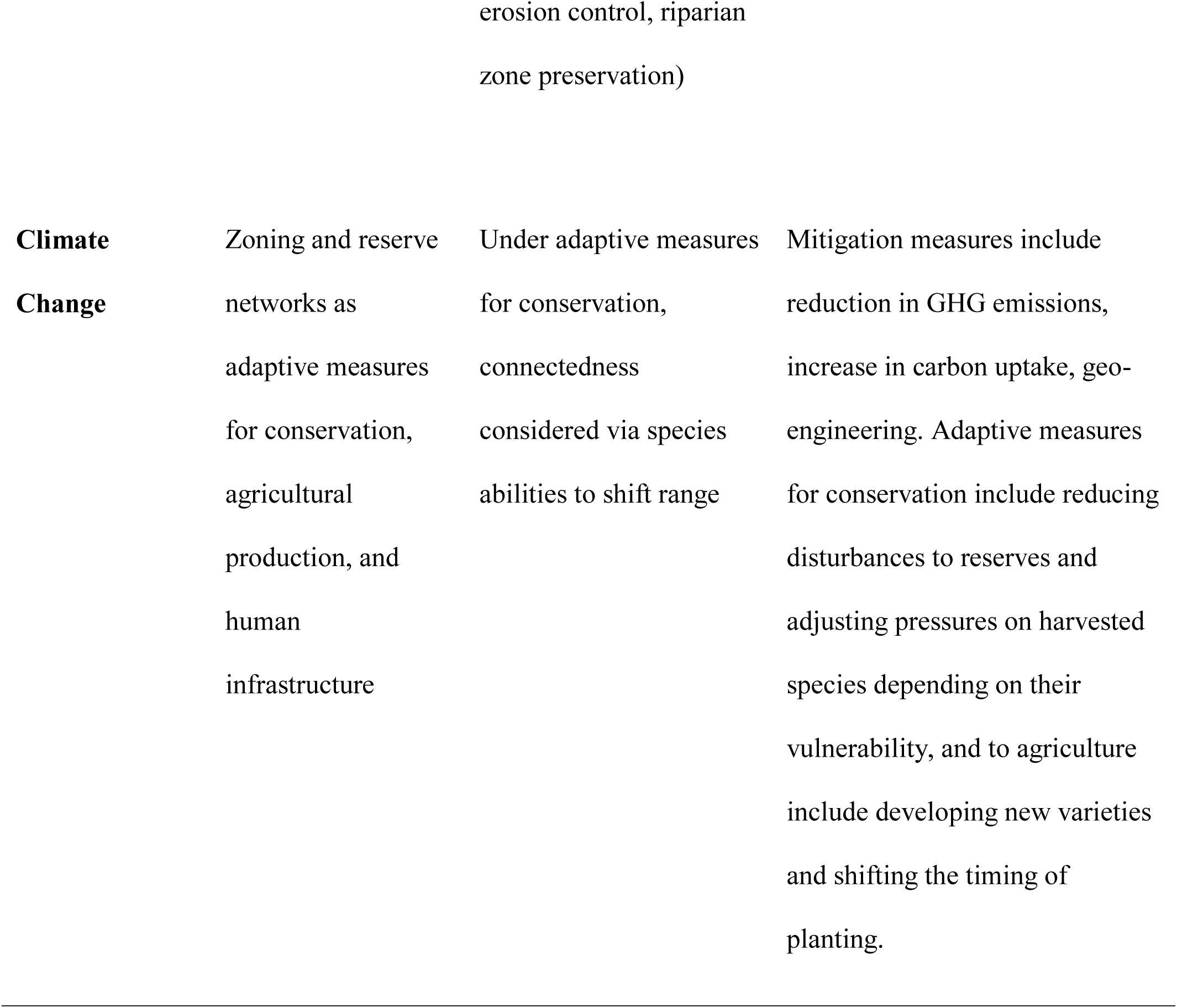
An overview of management actions taken by practitioners in response to drivers of environmental change (IUCN 2000, MEA 2005b,c,d,e,f, Grafton et al. 2010) organized under the framework of the key features of metaecology. Columns represent management actions directed at controlling the regional spatial structure of driver; its connectedness; and the driver, or its direct causes and consequences, at the local scale. Responses to climate change are divided into mitigation responses (directed at stabilizing climate change) and adaptive responses (directed at managing the risk of climate change).

Following rows in Table 2, one observes that elements of metaecology are present among current management actions for all drivers. However, it is also clear that there are important biases. Actions to mitigate overharvesting, pollution and climate change are strongly focused on local scale measures, even if these measures are to be replicated many times in space. In contrast, actions addressing biological invasions explicitly aim to control regional connectedness in order to prevent new introductions and stop the spread of invasive species. This approach is likely common because it is well-established that invasion prevention is more cost-effective than eradication or containment of invasive species (IUCN 2000, Lodge et al. 2016). Uniquely, actions to mitigate the consequences of land use change often invoke all three elements of metaecology. Land use zoning typically encompasses the spatial distribution of protected areas, a buffering region to reduce the impact of surrounding land uses, and corridors to facilitate connectivity (Lindenmeyer and Fischer 2006). Clearly, incorporation of unrepresented key features of metaecology could modify proposed actions or identify management alternatives for all drivers.

Following columns in Table 2, it becomes apparent that ecological systems are subject to multiple drivers of environmental change. Thus, applying metaecological approaches should ideally incorporate responses to multiple drivers simultaneously. Regional planning in terrestrial landscapes should simultaneously target land use change, range shifts caused by climate change, mitigation of non-point source pollution, and overharvesting. Management plans in freshwater systems should acknowledge the hierarchical spatial structure of river basins and the disruption in river connectedness caused by river damming. Finally, management plans in coastal systems should simultaneously target overfishing, biological invasions, and the spread of pollutants of both terrestrial and aquatic origins.

## Supplementary Materials: Literature Review

1. Aim. We first conducted a bibliometric analysis regarding the employment of the meta-ecological framework in the applied ecology literature in general.

Methods. We conducted a literature search in the Web of Science for all articles appearing in the applied ecology literature using the words metapopulation (or meta-population), metacommunity (or meta-community) or metaecosystem (or meta-ecosystem) in the title, topic or abstract (March 16, 2016). Selected journals included Biodiversity and Conservation, Biological Conservation, Biological Invasions, Conservation Biology, Ecological Applications, Global Change Biology, Journal of Applied Ecology, and Restoration Ecology.

2. Aim. We then conducted a bibliometric literature analysis to assess whether researchers investigating the most important drivers of environmental change – namely land use change, climate change, pollution, biological invasions and overharvesting (MEA 2005) – are employing the meta-ecological framework as judged by terminology, study design, and literature cited; and whether they could, based on a description of the research question and study design, benefit from using this framework.

Methods. We conducted literature searches in Scopus (March 16, 2016) for all articles published in the last 5 years (2011-2016) containing words relative to land use change (“land*use change” or “habitat loss”, 1780 references retrieved), climate change (“climate change”, truncated at the first 2000 references in each year), pollution (“pollut*, truncated at the first 2000 references in each year), biological invasions (“invasive species” or “biological invasions”, 6712 references), and overharvesting (“overharvest*” or “overexploit*”, 650 references) in the title, abstract or keywords.

We then randomized each reference list and took the first 20 publications that satisfied the following criteria: (i) the publication was an article in a peer-reviewed journal (ii) the study included a live and natural biotic component (iii) the study was empirical (iv) the driver of environmental change in question was a central research focus in the study and (v) the above - mentioned criteria could be satisfactorily assessed by reading the Abstract.

The 120 publications were then divided among six pairs of researchers; within each pair, one researcher served as the primary reader for data extraction and the other as a reviewer. Some papers upon closer examination were considered not to adhere to the abovementioned criteria and were excluded from the analysis.

Data extraction consisted of marking whether (i) the article mentioned the words metapopulation (or meta-population), metacommunity (or meta-community) or metaecosystem (or meta-ecosystem) (ii) the article cited any of the relevant meta-ecological literature (for this categorization all articles were examined by a single reader familiar with the meta-ecological literature, Pedro Peres-Neto) (iii) the article described a research scenario that could potentially benefit from a meta-ecological approach, that is, articles were considered to potentially benefit from metaecology when the study design comprised multiple locations and when the research question suggested that flows of individuals or materials among locations could be important in influencing local ecological entities (i.e. populations, communities or ecosystems) and (iv) the research design included any or all of the key features of the meta-ecological framework, namely spatial structure (i.e., if multiple locations were considered, whether in a hierarchical fashion or not), flows of individuals, energy or matter (i.e. whether directly measured through, e.g., mark-recapture studies, estimates of genetic divergence or isotopic analysis of elements; or indirectly assumed based on, e.g., geographic distances, measurement of dispersal traits or analysis of physico-chemical differences through time and space), and key abiotic and biotic processes at the local scale (i.e., whether the study considered demography or interactions with the abiotic environment or other species).

